# Enrichable consortia of microbial symbionts degrade macroalgal polysaccharides in *Kyphosus* fish

**DOI:** 10.1101/2023.11.28.568905

**Authors:** Aaron Oliver, Sheila Podell, Linda Wegley Kelly, Wesley J. Sparagon, Alvaro M. Plominsky, Robert S. Nelson, Lieve M. L. Laurens, Simona Augyte, Neil A. Sims, Craig E. Nelson, Eric E. Allen

## Abstract

Coastal herbivorous fishes consume macroalgae, which is then degraded by microbes along their digestive tract. However, there is scarce foundational genomic work on the microbiota that perform this degradation. This study explores the potential of *Kyphosus* gastrointestinal microbial symbionts to collaboratively degrade and ferment polysaccharides from red, green, and brown macroalgae through *in silico* study of carbohydrate-active enzyme and sulfatase sequences. Recovery of metagenome-assembled genomes (MAGs) reveals differences in enzymatic capabilities between the major microbial taxa in *Kyphosus* guts. The most versatile of the recovered MAGs were from the Bacteroidota phylum, whose MAGs house enzymes able to decompose a variety of algal polysaccharides. Unique enzymes and predicted degradative capacities of genomes from the *Bacillota* (genus *Vallitalea*) and *Verrucomicrobiota* (order Kiritimatiellales) suggest the potential for microbial transfer between marine sediment and *Kyphosus* digestive tracts. Few genomes contain the required enzymes to fully degrade any complex sulfated algal polysaccharide alone. The distribution of suitable enzymes between MAGs originating from different taxa, along with the widespread detection of signal peptides in candidate enzymes, is consistent with cooperative extracellular degradation of these carbohydrates. This study leverages genomic evidence to reveal an untapped diversity at the enzyme and strain level among *Kyphosus* symbionts and their contributions to macroalgae decomposition. Bioreactor enrichments provide a genomic foundation for degradative and fermentative processes central to translating the knowledge gained from this system to the aquaculture and bioenergy sectors.

**Importance:** Seaweed has long been considered a promising source of sustainable biomass for bioenergy and aquaculture feed, but scalable industrial methods for decomposing terrestrial compounds can struggle to break down seaweed polysaccharides efficiently due to their unique sulfated structures. Fish of the genus *Kyphosus* feed on seaweed by leveraging gastrointestinal bacteria to degrade algal polysaccharides into simple sugars. This study is the first to build genomes for these gastrointestinal bacteria to enhance our understanding of herbivorous fish digestion and fermentation of algal sugars. Investigations at the gene level identify *Kyphosus* guts as an untapped source of seaweed-degrading enzymes ripe for further characterization. These discoveries set the stage for future work incorporating marine enzymes and microbial communities in the industrial degradation of algal polysaccharides.

## Introduction

The *Kyphosus* genus of herbivorous fish, commonly referred to as nenue or rudderfish, graze primarily on macroalgae (1). *Kyphosus* fish serve important ecological roles by controlling algal cover in Indo-Pacific (2) and Caribbean coral reefs (3), thereby mediating coral-algal competition and overall coral growth and benthic community composition (4). Their diverse diet includes macroalgae from the three major taxonomic groups: Rhodophyta (red), Chlorophyta (green) and Ochrophyta (brown) (1). Polysaccharides constitute as much as 60% of macroalgal cells by weight (5) and serve roles in both cell structure and energy storage (6). The complex network of linkages in structural polysaccharides resist degradation from chemical and enzymatic stressors and serves as a physical defense mechanism for algal cells (7).

Algal polysaccharides differ from common polysaccharides found in land plants due to the addition of sulfate ester groups (8). Structural polysaccharides from red algae include agar, carrageenan, porphyran, and xylan, which all contain such sulfate groups (9). Brown algae contains the sulfated polysaccharide fucoidan for structure as well as unsulfated alginate as a storage polysaccharide (9). Green algae contain sulfated polysaccharides such as xylan and ulvan but also contain large amounts of unsulfated cellulose common in land plants (9). Algal polysaccharides are depolymerized primarily through the enzymatic activity of bacterial glycoside hydrolases (GHs) and polysaccharide lyases (PLs) (10), two classes of carbohydrate-active enzymes (CAZymes) (11). Sulfated polysaccharides are particularly recalcitrant to digestion because an additional enzyme class, the sulfatases, is necessary for complete degradation. Full enzyme pathways for the breakdown of various algal polysaccharides have been proposed (9,12) that include both required CAZyme and sulfatase activities. However, not all algal polysaccharides have well-defined degradation pathways or unique associated CAZymes that enable a high-level connection between gene presence and catabolized substrates. Likewise, sulfatase classes within the SulfAtlas database (13) are primarily classified based on evolutionary history rather than substrate specificity or enzymatic activity, so our ability to evaluate pathway completeness *in silico* is limited.

Once complex carbohydrates are broken into subunits by CAZymes and sulfatases, they are utilized by gut microbiota in fermentation reactions to produce short-chain fatty acids (SCFAs) (14). The SCFAs acetate, propanoate, and butyrate have been previously measured in high quantities in *Kyphosus* hindguts (15) and are utilized by the host fish for energy (16). Previous work has suggested correlations between SCFA profiles and bacterial composition (15), but there is no genomic work in algivorous fish pinpointing which microbiota contribute to host nutrition in this way and what pathways are utilized to produce these essential SCFAs.

Our overall understanding of the role of gut microbiota in digestion is still limited in most fishes (17), including *Kyphosus*, in part due to a focus on gut composition and diversity rather than function. The genetic study of *Kyphosus* gut symbionts has been limited to 16S rRNA (15,18) and metabolomic (18) investigations until the incorporation of shotgun metagenomics in a few recent studies (19,20). What functional profiling has been done in fish guts often relies on extrapolation from amplicon-based taxonomic distributions (21–24), and no study has yet generated a large collection of metagenome-assembled genomes (MAGs) from an algivorous fish gut. A *de novo* genomic investigation of *Kyphosus* symbionts has the potential to reveal degradative capacities that cannot be extrapolated from taxonomic lineage or relatedness to database representatives.

Discoveries from better studied human gut and terrestrial herbivore systems provide suggestions for how *Kyphosus* symbionts might gain and use such gene pathways. Human gut bacteria have acquired enzymes which degrade sulfated algal polysaccharides through horizontal gene transfer (25,26). Horizontal gene transfer of antibiotic resistance genes has also been observed in fish gut biofilms (27), but this phenomenon has not yet been reported for carbohydrate-active enzymes in any fish gut symbiont microbe. Once acquired, CAZymes and sulfatases potentially originating from one or multiple organisms may then decompose algal polysaccharides in complex, stepwise pathways. A cooperative division of labor strategy, in which partial breakdown products from one bacterial population serve as a degradative substrate for other bacteria in the community, has been proposed to occur in human gut microbiota (28) and has been suggested as a way to improve polysaccharide degradation in engineered communities (29). The degree to which collaboration may occur in the herbivorous fish gastrointestinal tract remains unknown.

Exploring functional diversity not only improves our understanding of herbivorous fish digestion but may also enable concrete applications in the fields of aquaculture and bioenergy. Most aquaculture is currently sustained through compound feeds that are composed of fishmeal and fish oils from wild-caught fish (30). Although innovations in aquaculture feed have lowered the trophic levels of captive carnivorous fish and improved overall feed efficiency (31), concerns about sustainability and food security remain. Wan *et al.* (2019) argue that the discovery of efficient methods to degrade complex polysaccharides and enhance nutrient digestibility is a key knowledge gap and barrier limiting macroalgae inclusion into commercial aquafeeds (32). Macroalgal feed additives are also known to counteract methanogenesis in terrestrial ruminants (33) and thus can be applied to reduce methane emissions from livestock husbandry. However, deficiencies in ruminant microbiome digestive capacities may influence the future development and long-term success of seaweed dietary supplementation strategies. Research on *Kyphosus* symbionts and their enzymes can inspire commercializable and scalable methods to break down these barriers in industry.

Innovations exploiting the experimental propagation of enrichment cultures with *Kyphosus* symbionts can harness these microbial communities for further study and experimentation with commercial outputs in the bioenergy sector as well as the development of macroalgal feed supplements. While a few bacterial isolates have been recovered and sequenced from kyphosid guts (34), no previous study has enriched entire communities from these fishes to investigate their hydrolytic and fermentative capabilities. Hydrolysis of carbohydrates, proteins, and lipids into their monomeric components is a key step in biogas and bioethanol production from macroalgae (35,36), and the degradation of algal polysaccharides is often the rate limiting step in anaerobic digestion (37). Milledge *et al.* (2019) call for future studies to look beyond commercially available enzymes to discover candidates that can more efficiently degrade algal polysaccharides (38). The *Kyphosus* gut, with its understudied functional diversity and degradative pathways, offers an untapped source of such enzyme and inoculum candidates.

This study leverages metagenome-assembled genomes from *Kyphosus vaigiensis, Kyphosus cinerascens, and Kyphosus hawaiiensis* gut symbionts and inoculated bioreactor enrichments to connect whole genome degradative potential of algal polysaccharides to accurate taxonomic lineages and functional roles. The addition of genomes from bioreactor enrichments explores leveraging the metabolic capacities of *Kyphosus* gut consortia in industrial processes. This work extends previous studies of taxonomic-level biogeography (18) and contig-level gene associations (15,20) in this system using high-quality MAGs, which enables differentiation between processes that can potentially be executed within a single cellular compartment (individual microbial species/population) and those likely to require cooperative action by multiple cells from different species (community impacts). Discoveries in this study provide foundation for genome-level understanding of microbial contributions to herbivorous fish digestion and beget future investigations to apply these findings towards applications in the aquaculture and bioenergy sectors.

## Materials and Methods

### Sample description and metagenomic assembly

DNA was extracted from liquid samples from ten anaerobic bioreactors inoculated with gut contents from two *Kyphosus* fishes (**Table S1**) using methods previously described (18) and propagated to enrich degradative properties. Samples were taken 9-10 days after inoculation and incubation at 30°C. Anoxic cultures of 50ml were processed in a portable anaerobic chamber containing one-third strength sterile artificial seawater (Instant Ocean, Spectrum Brands, Blacksburg, VA) in 150ml serum bottles, crimp sealed with a rubber septum. Approximately 1g of fish gut section contents were placed in the bottles along with the indicated substrate (**Table S1**) and sealed, with no additional feedstock added before sequencing.

Samples were sequenced using Illumina NovaSeq 6000 technology (Illumina, San Diego, CA). Read trimming was performed using Trimmomatic v. 0.36 (39) with the following parameters: adapter-read alignment settings 2:30:10, LEADING:10, TRAILING:20, HEADCROP:12, SLIDINGWINDOW:4:15, MINLEN:200. Taxonomic composition of metagenomic reads was determined using Kraken v. 2.0.9 (40), with taxonomic assignment using a protein database based on all amino acid sequences in the NCBI nr database (41) as of April 2022. Cleaned reads were assembled in metaSPAdes v. 3.13 (42) with a minimum contig retention size of 2000 nucleotides.

### Gene calling and functional annotation

Gene boundaries were predicted using prodigal v. 2.6.2 (43) and annotated using prokka v. 1.12 (44). Genes were assigned to CAZy classes from the dbCAN HMMdb v. 10 database (45) based on the CAZy database (11) and to sulfatases classes from the SulfAtlas v 2.3 database (13), using methods previously described (20). Signal peptides were identified using SignalP v. 6 (46) with default parameters.

Enzyme novelty was evaluated using DIAMOND blastp (47) searches against the NCBI nr database (41) as of April 2022. Some CAZyme classes were grouped into the category of “peptidoglycanases” using the division proposed by López-Mondéjar *et al.* (2022) (48). Distributions of annotated proteins were compared to free-living relatives from the OceanDNA database (49).

### Metagenomic binning and biosynthetic gene cluster prediction

Metagenomic binning was performed using MetaWRAP v. 1.3.2 (50) with a minimum completeness cutoff of 0.7 and a maximum contamination cutoff of 0.05 as determined by CheckM v. 1.0.12 (51). MAG taxonomy was determined using GTDB-Tk v. 1.5.1 (52) with release 202 of the Genome Taxonomy Database (53).

Viral contigs and prophage were identified using DeepVirFinder v. 1.0 (54) using a q-score cutoff of 0.94. Viral sequence completeness was determined using Checkv v. 1.5 (55), we only retained regions marked as “high-quality” or “complete”. Viral sequences were assigned host taxonomies using VPF-class (56).

Biosynthetic gene clusters (BGCs) were predicted for each MAG using antiSMASH v. 6.1 (57). Predicted products and BGC classes were annotated using BiG-SLiCE v. 1.1.1 (58). Gene cluster distances were calculated using the BiG-FAM webservice v. 1.0.0 (59), using a novelty distance cutoff of 900 following previous studies (59–61). Short chain fatty acid gene clusters were annotated using gutSMASH v. 5.0.0 (62).

### Phylogenomics and enzyme phylogenetics

A phylogenetic tree of MAGs was generated using PhyloPhlAn v. 3.0.2 (63) using a concatenated universal set of 400 marker genes (64). MAGs containing at least 100 marker genes underwent concatenated alignment using mafft v. 7.505 (65). The phylogenetic tree was built using RaxML v. 8.2.12 (66) and visualized using R v. 4.2.0 (67) packages treeio v. 1.20.0 (68), ggtree v. 3.4.0 (69), and ggtreeExtra v. 1.6.0 (70).

Multiple sequence alignments for genes belonging to CAZy class GH86 were made using MUSCLE v 3.8.31 (71) and visualized using the R package ggmsa v. 1.2.0 (72). Gene trees were created using FastTree v. 2.1.10 (73). Additional reference genes were included in the tree based on DIAMOND blastp hits to the NCBI nr database as of April 2022. Protein domains were analyzed with the CDD webservice (74). 3D protein structures for CAZymes were predicted using ColabFold v. 1.3.0 (75) and visualized using ChimeraX v. 1.3 (76). Residue conservation was visualized using the WebLogo (77) webservice.

### Data availability

All custom code used for data analysis and visualization are available at https://github.com/AaronAOliver/KyphosusMAGs. Sequence reads are available under SRA bioproject numbers PRJNA819194 and PRJNA1023379. Complete MAG sequences and predicted proteins are available on Zenodo (https://zenodo.org) under DOI no. 10.5281/zenodo.8277654.

## Results

### A (meta)genome catalog of enrichable symbionts in the *Kyphosus* gut

New data derived from *K. cinerascens* and *K. hawaiiensis* enrichment cultures expands the diversity of previous *K. cinerascens*, *K. hawaiiensis*, and *K. vaigiensis* gut metagenomes (20). This more complete catalog of *Kyphosus* gut microbiota provides additional details on the metabolic potential of taxa that were rare in the *in vivo* gut metagenome samples and highlights potential challenges in harnessing gastrointestinal microbiota for industrial processes. The fish inoculum species, gut location, and feedstock that were combined to establish each enrichment sample are described in **Table S1**. The taxonomic classification of unassembled metagenomic reads revealed a surprising consistency between the *in vivo* gut microbiomes (20) and enrichment samples (**Figure 1**). *Bacillota*, *Bacteroidota*, and *Gammaproteobacteria* constitute the dominant bacterial lineages in most samples, although the *Desulfovibrionales* order (phylum *Thermodesulfobacteriota*) was highly abundant in two enrichment samples.

**Figure 1.**
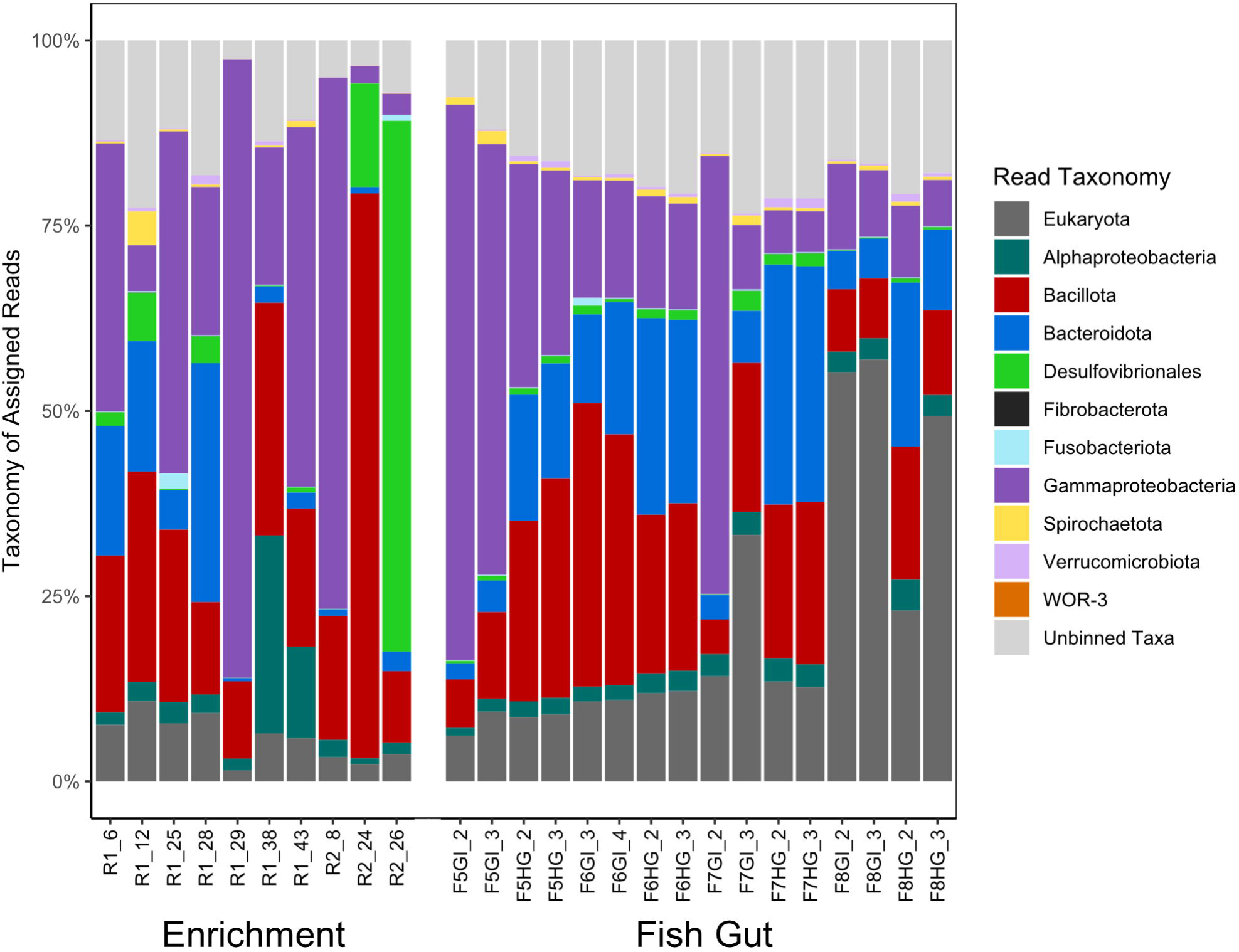
Taxonomic distribution of enrichment and fish gut samples. The taxonomic distribution of unassembled classified reads as determined using Kraken2. Any taxonomic lineages that are not associated with a binned MAG are grouped into “unbinned taxa.”

211 medium and high-quality MAGs were binned from the *in vivo* fish gut metagenomes and newly assembled enrichment metagenomes. These MAGs all met the minimum 70% completion, maximum 5% redundancy standards (78). The number of recovered MAGs per metagenome is shown in **Figure S1**. The assembly statistics for enrichment metagenomes are shown in **Table S2** and MIMAG-compliant (78) summary information are shown in **Table S3.** Consistent with the unassembled read-based taxonomic profiles of the metagenomes, most MAGs were assigned to the phyla *Bacillota* (78 MAGs), *Bacteroidota* (72), the class *Gammaproteobacteria* (31), the class *Desulfovibrionales* (13), or *Verrucomicrobiota* (6). The enrichments provide access to data on microbial members that were not as abundant in the fish gut metagenomes and vice versa. In one example, the *Verrucomicrobiota* class *Kiritimatiellales* was binned in fish gut samples but not in enrichment metagenomes. This novelty was reflected in nucleotide similarities, as only 9 of the 74 (12%) enrichment MAGs match MAGs generated from *in vivo* fish gut metagenomes at the species level.

Viral sequences comprised less than 0.5% of all unassembled metagenomic reads, with 69 viral contigs and 3 prophages identified as either high quality or complete. With these viral elements, 30 auxiliary metabolic genes found on potential prophage regions were annotated as CAZymes, and 13 as sulfatases, suggesting a potential role for viral dissemination of these genes across the bacterial community. The taxa *Bacillota*, *Bacteroidota*, and *Gammaproteobacteria* were the most frequently predicted viral hosts, which is consistent with the taxonomic abundances of classified unassembled metagenomic reads and recovered MAGs (**Table S4**). Despite the presence of numerous auxiliary metabolic genes generally related to polysaccharide degradation, none of the viral sequences we detected appeared to specifically target large, complex sulfated macroalgal polysaccharides.

### Genome capacities reveal metabolic specialization among gut symbionts of *Kyphosus* fish

The distribution of CAZymes and sulfatases was correlated with the phylogeny of fish gut and enrichment MAGs (as determined through a concatenated marker gene tree, **Figure 2a**). This assessment revealed that among the MAGs generated in this study, the *Bacteroidota* genomes contained the majority of CAZymes and sulfatases (**Figure 2b**). Algal degradation-specific CAZyme-rich genomes among the MAGs from other phyla were restricted either to a single order, *Kiritimatiellales* (*Verrucomicrobiota*), or a single genus, *Vallitalea* (*Bacillota*). Recovered *Gammaproteobacteria* and *Desulfovibrionales* genomes lacked enzymes required for digesting sulfated algal polysaccharides, despite the relatively high abundance of these taxonomic groups in classified unassembled reads and the recovered MAGs. However, the *Gammaproteobacteria* MAGs contained more peptidoglycanases than other taxa, suggesting a niche in digesting alternative dietary components. This analysis also showed that CAZymes targeting ulvan, a green algal polysaccharide, were less prevalent among the obtained symbiotic MAGs than CAZymes targeting red and brown algae-associated polysaccharides (**Figure 2b**), consistent with previous results quantifying relative amounts of these algae types consumed by the *Kyphosus* fish included in this study (20). The most abundant phyla all had binned MAGs from both *in vivo* and enrichment samples (**Figure 2c**).

**Figure 2.**
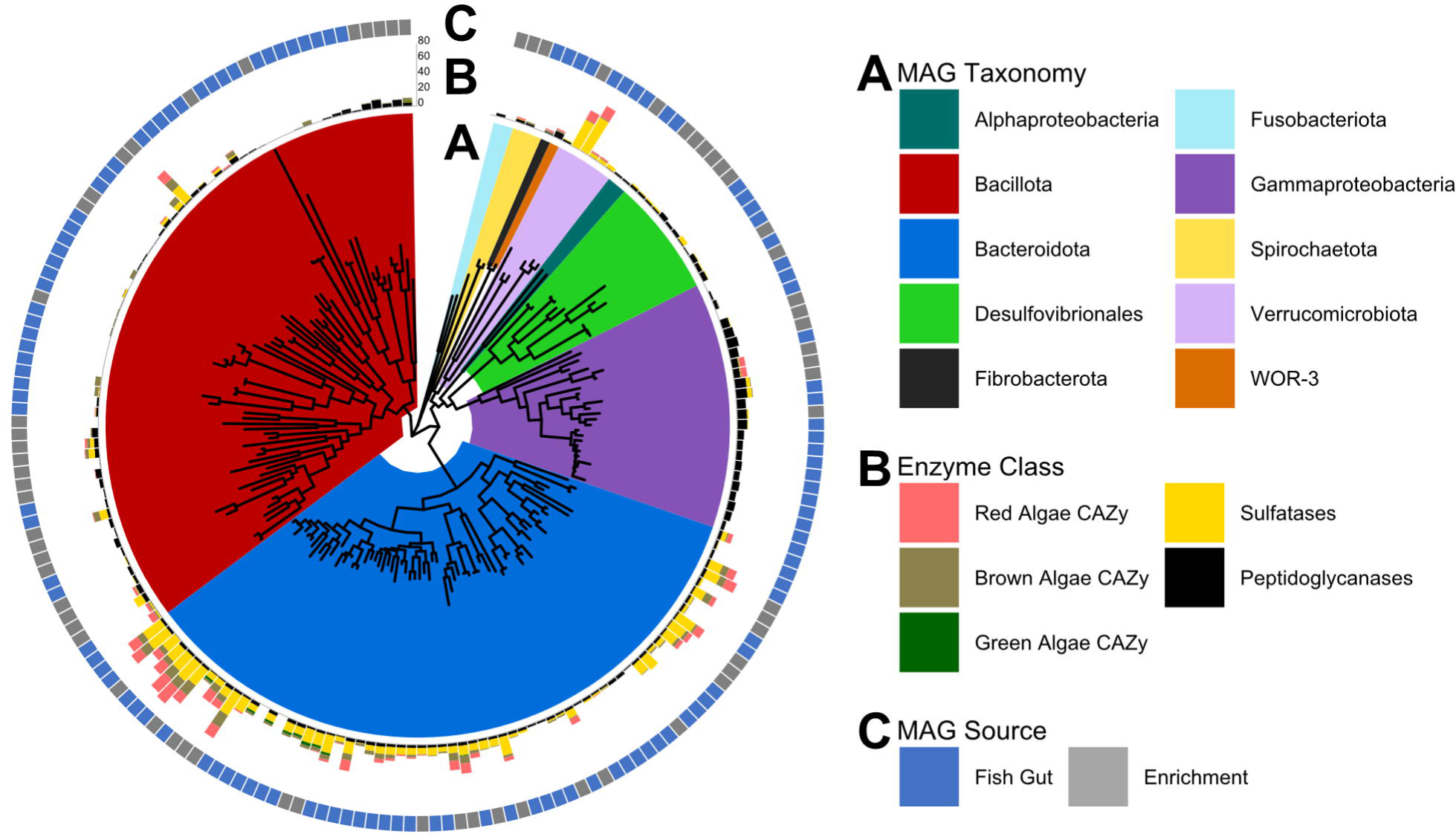
Genomic CAZyme distributions reveal connections between metabolic strategies and taxonomic lineage. (A) The gene tree shows a concatenated alignment of 400 PhyloPhlAn universal marker genes for each recovered MAG, with branches colored by assigned MAG taxonomy. (B) The inner ring displays genomic gene counts for sulfatases and carbohydrate-active enzymes that specifically target algal polysaccharides or peptidoglycan. (C) Environmental source of each MAG.

An assessment of SCFA production gene pathways of recovered MAGs using gutSMASH (62) revealed that most of the *Kyphosus* gut symbiotic taxa (67% of fish gut MAGs, 77% of enrichment MAGs) can potentially contribute to host nutrition through the production of SCFAs (**Figure 3**). 139 genomes from analyzed kyphosid fish gut microbial communities contained pathways for producing acetate but only six genomes contained pathways for butyrate production. The pyruvate formate lyase and pyruvate:ferredoxin oxidoreductase pathways were the most abundant overall, present in 126 MAGs, while *Bacteroidota* contained the most gene clusters (39) related to propanoate production.

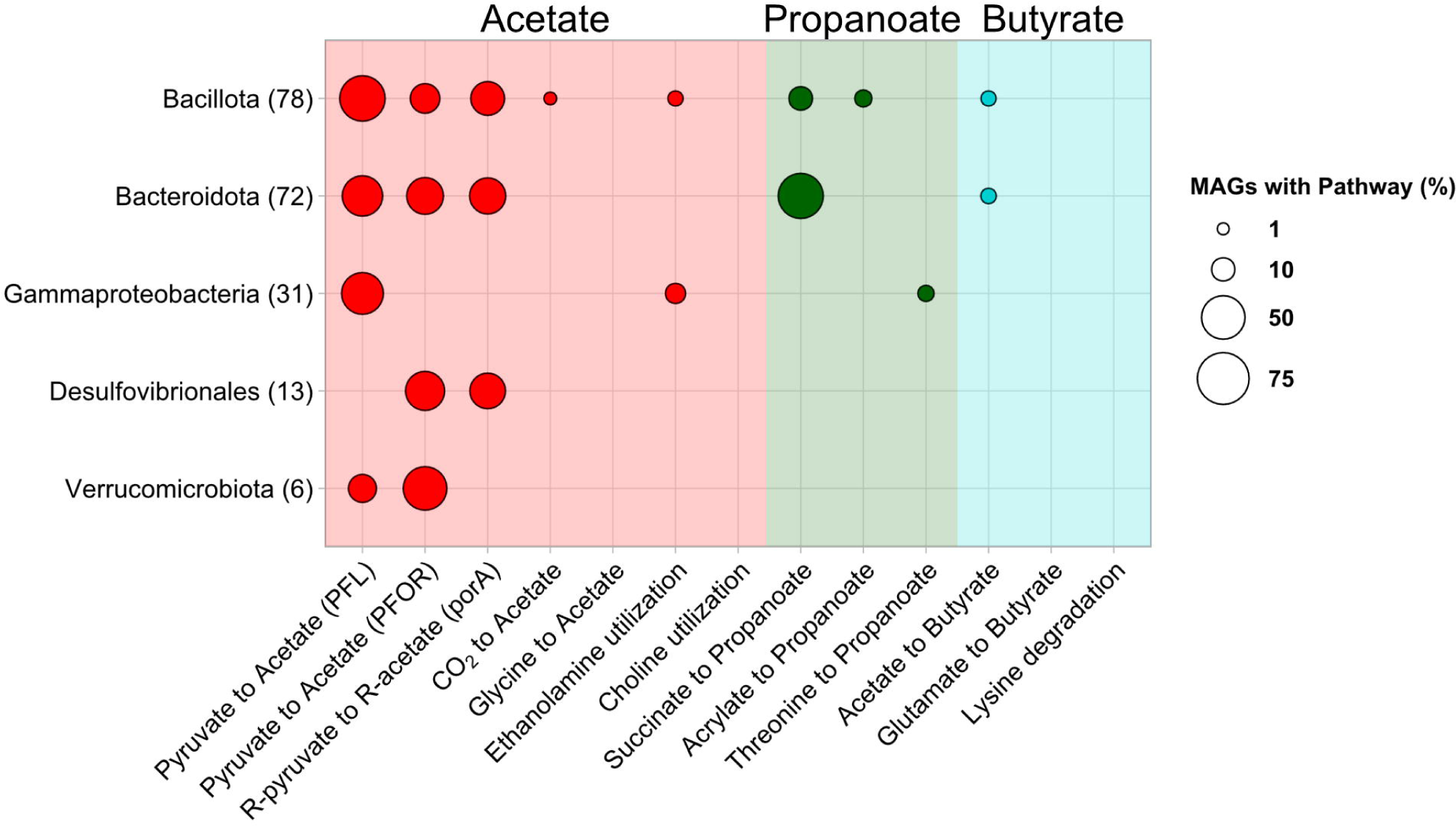

The overall prevalence of acetate pathways was lower than that found previously in human gut microbiota. The total absence of some alternate fermentation pathways from our MAGs, such as choline utilization, suggests that those processes are not core to dominant members of the *Kyphosus* gut microbiome. Only one genome from this study contained fermentation pathways involving the degradation of amino acids such as glycine, threonine, and lysine, suggesting that *Kyphosus* gut microbiota do not rely directly on dietary proteins for energy. Such lessened reliance on nitrogen-based substrates for fermentation is consistent with a low protein, algae-based diet rich in available polysaccharides and limited in available nitrogen.

### Functional adaptations to life in the *Kyphosus* gut

Adaptations to environmental conditions in herbivorous fish guts studied here are reflected in the high abundance of CAZyme classes specifically targeting algal polysaccharides. **Figure 4a** shows that the amino acid sequences of selected CAZyme classes abundant in our assembled metagenomes are well conserved across *Kyphosus* gut symbiont genomes. However, such enzymes are poorly represented in both specialty and general databases of previously described sequences, with closest enzyme homologs averaging below 60% sequence similarity for most of the examined CAZyme classes. Similar trends are observed for the sulfatase subclasses in these *Kyphosus* gut symbiont genomes (**Figure 4b**). Both cases denote the extent that this study expands known sequence diversity within these enzyme classes, potentially suggests new subclasses, and highlights unusual domains that may not be captured by current databases.

**Figure 4.**
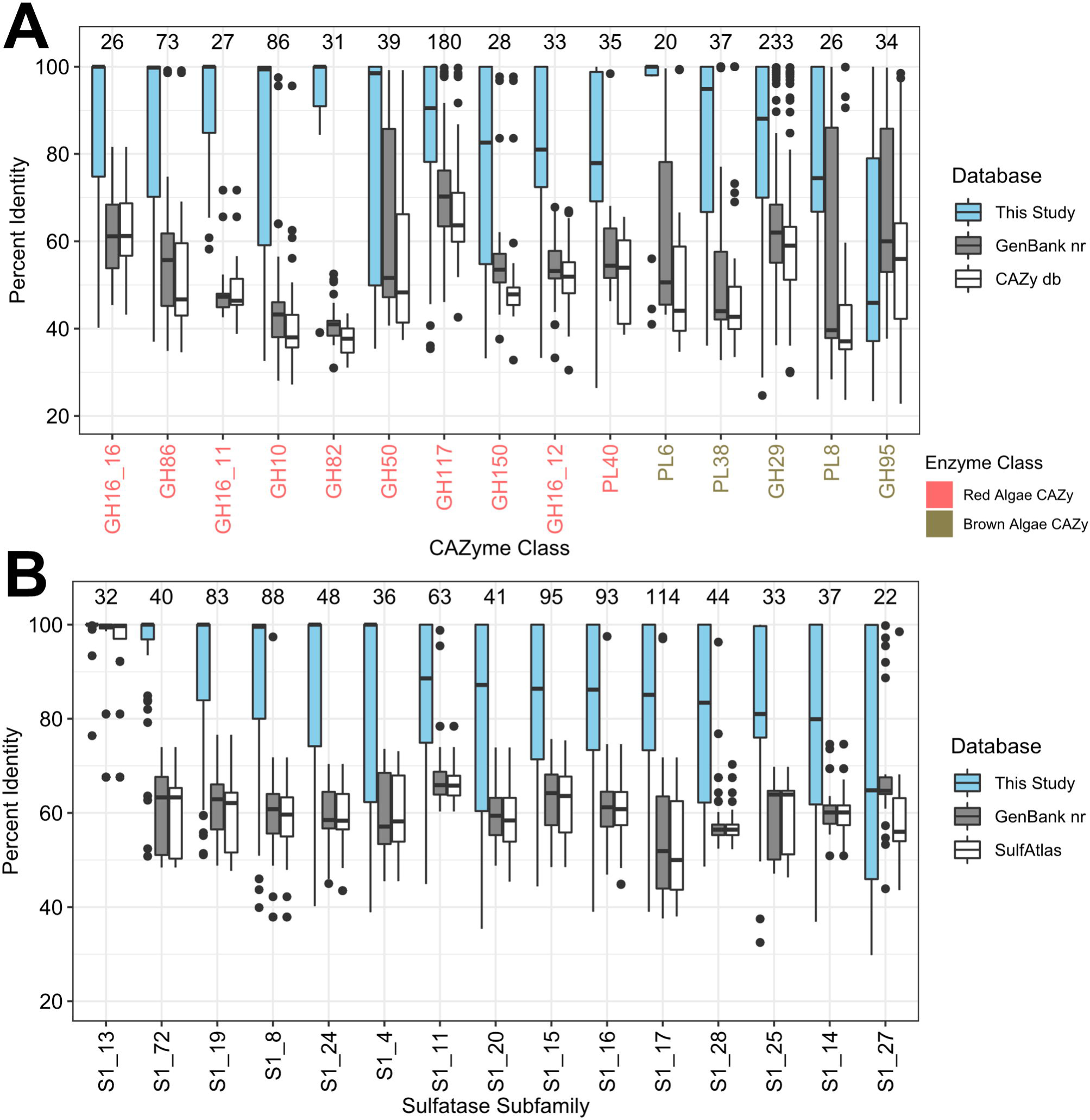
*Kyphosus* gut symbionts encode CAZymes and sulfatases divergent from other datasets and environments. Percent identity of binned (A) CAZymes and (B) sulfatases to best blast matches found in the following databases: all genes from MAGs in this study (orange), the GenBank nr database (green), and either (A) the CAZy database or (B) the SulfAtlas database (blue). Each group is labeled by the number of genes with that enzyme annotation found in our MAGs.

The addition of novel enzymes sequences to each of these enzyme classes presents numerous opportunities to expand our understanding of marine polysaccharide degradation. One example using the phylogeny of CAZy class GH86, consisting of β-agarases and β-porphyranases, illustrates previously unappreciated cryptic variability within this enzyme family. A gene tree of class GH86 CAZyme examples from this study (**Figure 5**), that includes the closest GenBank homologs, shows that most of the genes are associated with *Bacteroidota* from

**Figure 5.**
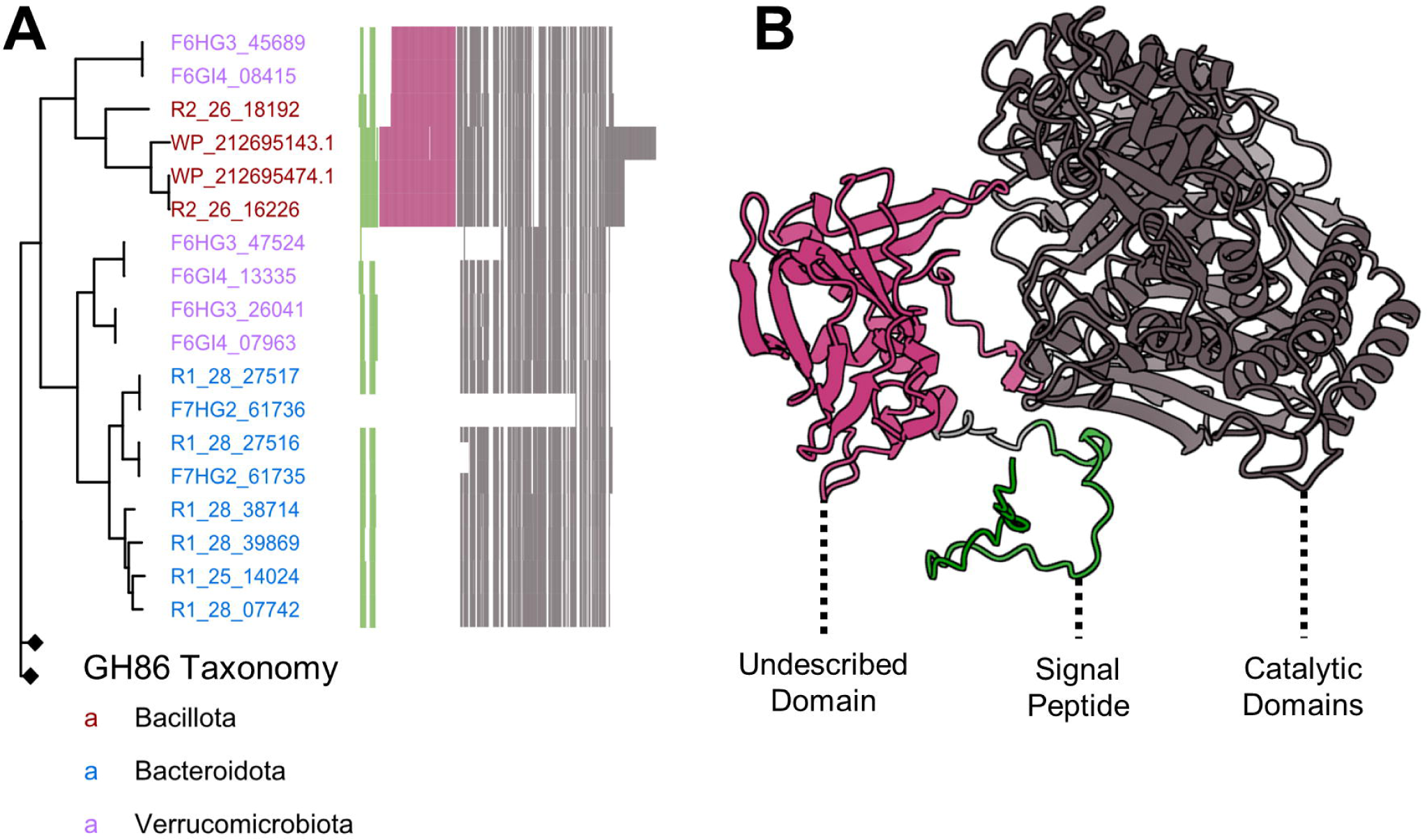
A β-agarase/β-porphyranase gene tree highlights an undescribed protein domain present in multiple phyla. (A) A gene tree of binned GH86 enzymes, with gene names colored by genome taxonomy. Nodes with black diamonds represent collapsed clades without the undescribed domain. A multiple sequence alignment is appended to the end of the tree, with colored vertical lines representing amino acid positions and white vertical lines representing gaps. (B) The predicted protein structure of GH86 enzyme R2_26_16226, with conserved CAZy domains highlighted in gray, the predicted signal peptide in green, and the conserved undescribed domain in pink. An uncollapsed version of the gene tree is included as **Figure S2** and a motif logo of the domain is included as **Figure S3**.

*Kyphosus* guts. This is consistent with the high abundance of CAZymes and sulfatases found among MAGS from the phylum (**Figure 2)**. Surprisingly, two GH86 genes recovered in *Bacillota* MAGs from bioreactor enrichments and two homologs from the NCBI nr database cluster together with two genes found among hindgut MAGS from the phylum *Verrucomicrobiota*, suggesting potential horizontal gene transfer from marine sediment communities into *Kyphosus* gut microbiota. Binned genes annotated as β-porphyranases all originate from hindgut or enrichment samples, consistent with previously reported physiological localization of polysaccharide degradation capabilities (20).

The multiple sequence alignment in **Figure 5a** highlights a unique pattern within the Bacillota genus *Vallitalea* and neighboring *Verrucomicrobiota* CAZyme sequences that has not been described in prior literature. This pattern might either extend the signal peptide or add an additional uncharacterized domain between the signal peptide and the porphyranase catalytic subdomain (79). Among NCBI nr homologs, only genes from an isolated *Vallitalea* genome (WP_212695143.1, WP_212695474.1) (80) contained this pattern. No other proteins in the GenBank nr database contained sequences matching this region at greater than 50% amino acid identity for this pattern of approximately 168 amino acids, with few conserved residues among our sequenced examples (**Figure S3**). Outside of the clade containing this novel domain, variability occurs primarily in the putative signal peptide region at the N-terminus of the protein, while the porphyranase domain itself is far more conserved. **Figure 5b** displays the predicted 3-dimensional structure of a *Kyphosus* symbiont GH86 enzyme, with the additional uncharacterized structure positioned between the predicted signal peptide and annotated catalytic β-agarase and β-porphyranase domains. This uncharacterized domain might influence an array of enzymatic properties such as a novel substrate specificity, concentration dependence, improved efficiency, or tolerance of different abiotic conditions. Although the function of this domain cannot be determined bioinformatically, this example is an interesting candidate for further enzymatic characterization and shows the promise of uncovering novel enzyme activity within the metabolic repertoire of the *Kyphosus* gut.

MAG sequences were interrogated using antiSMASH biosynthetic gene cluster detection software to determine whether *Kyphosus* gut-associated microbial taxa might encode any unusual secondary metabolites. The majority of *Bacillota*, *Bacteroidota*, *Verrucomicrobiota*, and *Gammaproteobacteria* MAGs from both fish gut inocula and bioreactor enrichments encoded BGCs typical of taxonomic relatives found in other vertebrate gut environments, such as lanthipeptides, betalactone, and arylpolyene (81,82). However, BGCs were not particularly abundant in our MAG catalog relative to other similar genomes. Our recovered *Gammaproteobacteria*, *Bacillota*, and *Bacteroidota* average fewer BGCs per genome than a random set of seawater MAGs of each taxonomic group from the OceanDNA database. Thus, our host-associated MAGs contain fewer BGCs per genome than their free-living relatives.

A total of 307 BGCs were annotated within our MAGs (**Figure 6**). 23% of annotated BGCs were determined to be complete based on BiG-FAM. 20 BGCs represent putative novel gene cluster families as determined by BiG-FAM (**Figure 6b**). These novel gene cluster families may represent unique natural products or enzymes specialized to the *Kyphosus* gut environment. Complete biosynthetic gene cluster annotations, novelty assessment, and associated taxonomy are included in **Table S5**.

**Figure 6.**
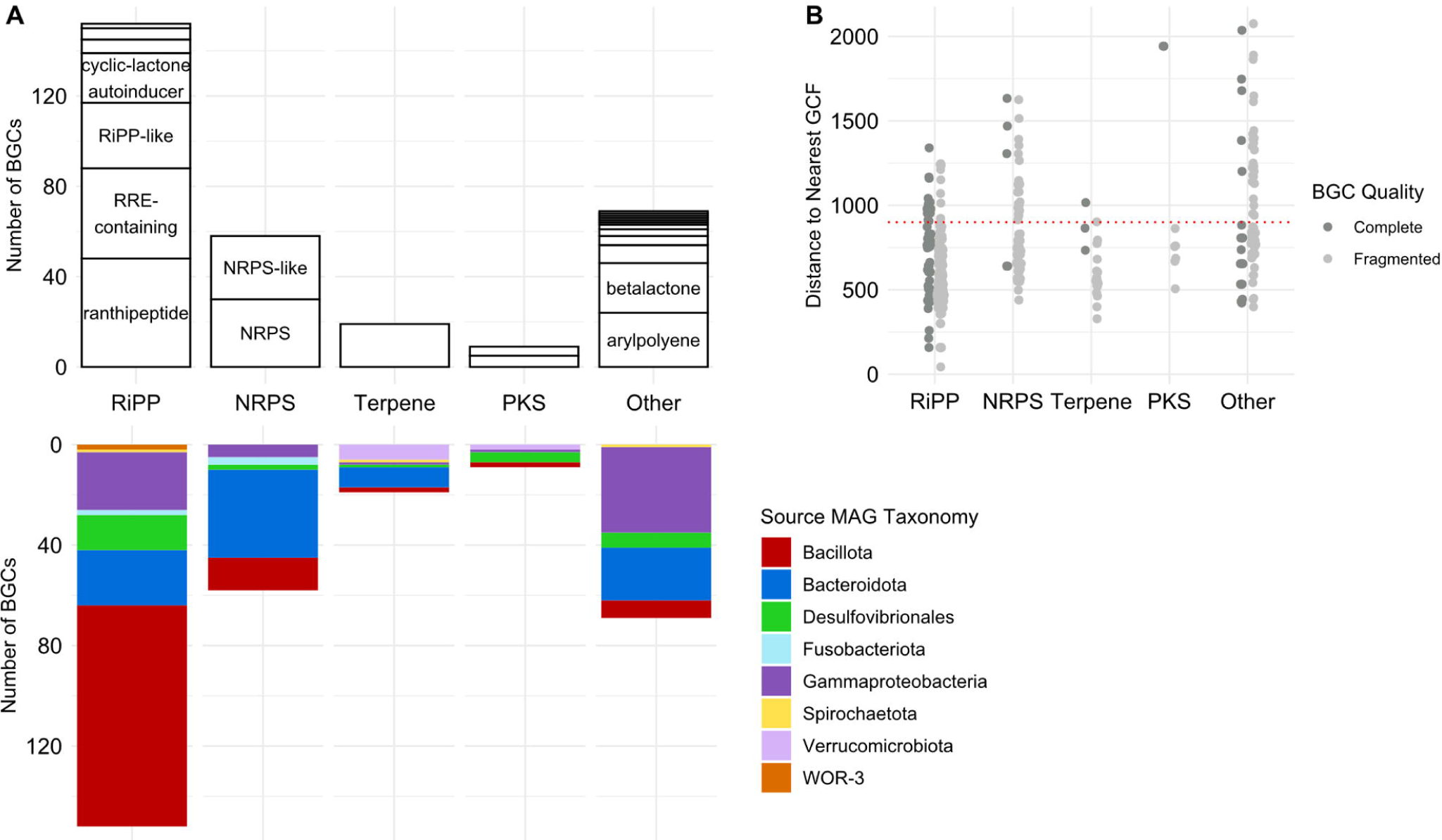
*Kyphosus* gut symbiont MAGs encode novel biosynthetic gene clusters. (A) On the positive y-axis, counts of binned BGCs grouped by BiG-SLiCE class and labeled by predicted product. On the negative y-axis, counts of binned BGCs grouped by BiG-SLiCE class and colored by associated MAG taxonomy. (B) Distance of binned BGCs to the nearest gene cluster family as determined by BiG-FAM. A distance above 900, marked by a dashed red line, suggests novelty and divergence from previously described gene cluster families. BGCs are colored orange if they are annotated as complete by BiG-FAM. Abbreviations used: RiPP, ribosomally synthesized and post-translationally modified peptides; RRE, RiPP recognition element; NRPS, non-ribosomal peptide synthetase; PKS, polyketide synthase.

### Community digestion of complex algal polysaccharides

Polysaccharide digestive capabilities vary among MAGs from different microbial taxa in the *Kyphosus* fish gut community, as shown in **Figure 7**. Despite overall microbiome-wide diversity, the MAGs generated in this study show that few individual genomes contain all of the enzymes necessary to completely degrade even a single type of complex algal polysaccharide, let alone the huge variety of natural variants characteristic of marine macroalgae (83) that might be ingested by generalist herbivorous fishes. Each microbial genome instead contains a limited assortment of enzymes capable of partially degrading a selection of different carbohydrate moieties, including potentially incomplete breakdown products generated by other microbes. Thus, combined pangenomic capabilities of several taxonomic groups appear to contain complementary collections of exported CAZymes that might facilitate adaptation to unpredictable variability in available polysaccharide content.

**Figure 7.**
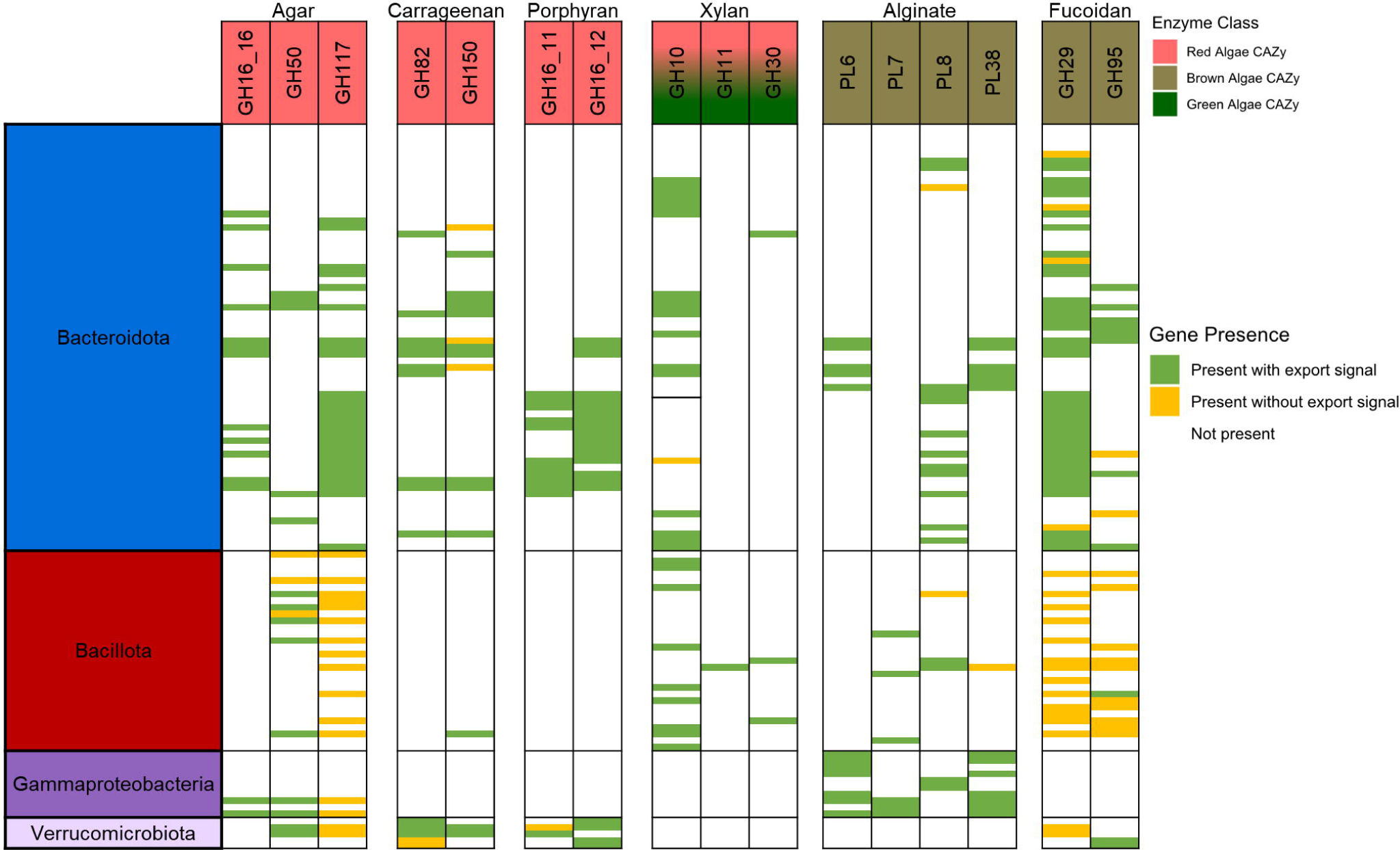
*Kyphosus* gut symbiont MAGs encode the capacity to degrade various algal polysaccharides collaboratively, but not solitarily. Each row represents a single MAG from the annotated taxonomic lineage. Only MAGs from the four lineages with the highest concentration of CAZymes (Bacillota, Bacteroidota, Gammaproteobacteria, and Verrucomicrobiota) are shown. MAGs with no applicable CAZy classes are not shown, and CAZy classes not associated with a single substrate or not found in any MAG are not shown. Green bars denote a signal peptide annotated to at least one of the appropriate CAZyme in a single MAG, while yellow bars mark the absence of a signal peptide on all appropriate CAZyme candidates within a MAG.

Levels of contribution to community-wide degradation of algal polysaccharides through extracellular enzymes are dependent on both cell taxonomy and targeted substrate. More than 90% of CAZymes that target macroalgal polysaccharides from *Bacteroidota* MAGs contain signal peptides that indicate export or integration into the cellular membrane. CAZymes in *Bacillota* MAGs largely lack these signal peptides in enzymes predicted to degrade fucoidan and agar, but the signal peptides are more abundant in the smaller set of CAZymes targeting xylan and alginates. Few *Bacillota* MAGs contain all the enzymes required to fully degrade complex algal polysaccharides such as porphyran, suggesting that cells from this taxonomic group might scavenge partial breakdown products degraded extracellularly by other taxa.

*Verrucomicrobiota* polysaccharide digestion enzymes appear to be more specialized towards red algae, with genomes consistently containing CAZymes predicted to digest agar, carrageenan, and porphyran. However, MAGs from this phylum seem to be lacking enzymes predicted to target green or brown algal polysaccharides. *Gammaproteobacteria* MAGs appear to have more enzymes involved in the digestion of non-sulfated polysaccharides such as alginate, and occasionally enzymes involved in agar degradation. Thus, the *Gammaproteobacteria* symbionts analyzed here have likely specialized in polysaccharide types that are easier to digest.

## Discussion

The recovery and characterization of 211 MAGs from *Kyphosus* gut and enrichment metagenomes connect detailed taxonomic classification with the potential of the major microbial contributors to digest complex algal polysaccharides. Algal polysaccharide-targeting enzymes from this study are divergent in sequence from previously sequenced and characterized representatives from other environments, clarifying prior assumptions about the metabolic capacities of this system using 16S rRNA or community composition. This work confirms and expands earlier work showing that certain members of the *Bacillota* and *Verrucomicrobiota* lineages are unexpectedly richer in some CAZyme and sulfatase enzyme classes than their respective taxonomic relatives (20). Differences between source inocula and the metagenomes of bioreactor enrichments inoculated with *Kyphosus* gut bacteria highlight potential challenges in harnessing these microbiota for bioenergy preprocessing of macroalgal feedstocks.

This study is the first to describe specific genes encoding SCFA production pathways in the genomes of fish gut microbiota. Microbial fatty acids serve as a key metabolite in gut-brain communication (84) and are a major source of available carbon for the host (85). SCFA pathway diversity is unexpectedly low for a system previously shown to contain high SCFA concentrations *in vivo* (16). However, this observation is consistent with a few dominant lineages, primarily the *Bacteroidota*, producing high amounts of SCFAs from the breakdown products of algal polysaccharides. Prior chemical work has observed that propanoate is more abundant than butyrate in *Kyphosus* guts (16), and our pathway enzyme abundance information at the genome level supports these observations (**Figure 3**). Likewise, observations in that same work noted rates of sulfate reduction were higher than methanogenesis, although both processes were negligible compared to SCFA production. This aligns with the low abundance of *Desulfovibrionales* and the near complete absence of Archaea in our metagenomes, which is consistent with repeated observations that dietary red macroalgae inhibit methanogenesis and thus the success of gut Archaea (33). Both sulfate reduction and methanogenesis appear to be minor sources of energy available for Kyphosid host absorption, compared to fermentation by *Bacteroidota* and *Bacillota*.

Herbivorous fish frequently contain visible amounts of sediment in their guts (86), which is thought to increase physical abrasion of gut contents and aid with degradation. Previous works have shown functional redundancy between the metabolic capacities fish gut and sediment communities (87), including carbon cycling. Gene flow has also been observed from fish feces to sediment microbiomes (88). Although the relative abundance of sediment-associated microbes in kyphosid fish microbiomes is low (20), one explanation for similar CAZymes and sulfatases between fish gut and sediment microbes could involve a circular loop of gene flow from fish guts to sediment through fecal pellets, and from sediments into fish guts through digestion. This hypothesis is supported by the fact that Kyphosid gut community *Bacillota* from the genus *Vallitalea* appear more closely related to marine sediment bacteria (80,89) than any previously reported examples from seawater or terrestrial gut microbiota (**Table S3**). Likewise, sediment-dwelling *Verrucomicrobiota* from the order *Kiritimatiellales* similar to those in *Kyphosus* fish guts have also been shown to degrade sulfated macroalgal polysaccharides (90), with genomes rich in both glycoside hydrolases and sulfatases (91). It is possible that consumption of sediment by *Kyphosus* fish improves polysaccharide digestion not only through physical breakdown of seaweed, but also by the contribution of additional enzyme capabilities originally derived from sediment bacteria that likely encounter highly diverse recalcitrant organic substrates including macroalgae biomass (92).

Although *Kiritimatiellales* MAGs recovered from fish guts contain more enzymes targeting algal polysaccharides than other members of their phyla, these taxa were not enriched in or recovered from enrichment metagenomes. This should not be problematic for enrichment processing if the dominant *Bacteroidota* contain CAZymes with overlapping specificities for the same substrates, as suggested in **Figure 7**. However future work will be needed to characterize detailed, sample-specific polysaccharide degradative chemistry using such a framework. *Vallitalea* and *Verrucomicrobiota* enzymes may also have some unique functionalities, as suggested by the extra domain present in their β-porphyranase sequences. Isolation and *in vitro* characterization of bioinformatically predicted enzyme activities will be necessary to integrate these discoveries into aquaculture and bioenergy applications.

Metagenomic data from the MAGs in this study suggest that few individual cells have the genomic potential to independently degrade all of the complex sulfated polysaccharide substrates present in marine macroalgae. However, secreted and extracellularly exposed transmembrane CAZymes may enable collaborative interactions between fish gut microbes to facilitate complete digestion of these molecules, without the high metabolic cost of encoding a complete, independent repertoire in every genome. A division of labor strategy cannot be fully confirmed without *in vitro* tests (93), although the first condition of functional complementarity appears to hold true between *Kyphosus* symbionts based on our bioinformatic investigations. In one similar study, gene based observations of complementarity for marine lignocellulose-degrading bacteria align with *in vitro* observations that support a division of labor hypothesis (94). Future work involving cultured representatives and enriched microcosms will be required to pin down the ecological strategies used by symbionts in this system.

This study provides a new baseline for *Kyphosus* microbiota at the genome level but begets a slew of new questions that require additional experimentation. Further work that connects enrichment composition, feedstock polysaccharide composition, and physical configuration to chemical measurements of degraded polysaccharides would help determine which phyla are required for complete polysaccharide breakdown. Isolation and characterization of divergent proteins with completely novel domains will determine what new enzymatic properties are unique to this system. Metatranscriptomic analyses utilizing the genome catalogs presented here will enable detailed analysis of substrate-specific metabolic pathway expression and species collaboration. *Kyphosus* digestive systems have long been studied as models for herbivorous fish gut fermentation and can now be explored further using these additional techniques to deliver a deeper understanding of their degradative and fermentative capabilities.

## Conclusion

Among the first metagenome-assembled genomes recovered from herbivorous fish guts and corresponding bioreactors, a new genomic catalog of *Kyphosus* gut symbionts highlights untapped diversity in enzymatic and collaborative potential in the degradation of algal polysaccharides. The enzymes encoded within these symbiont genomes are divergent from the extent of sequenced CAZymes, supporting the promise of herbivorous fish guts as a source of novel and industrially relevant enzymes. Expansion of these discoveries will not only clarify ecological interactions but have the potential to improve the applicability of macroalgae in the bioenergy and aquaculture sectors.

## Acknowledgements

This publication includes data generated at the UC San Diego IGM Genomics Center utilizing an Illumina NovaSeq 6000 that was purchased with funding from a National Institutes of Health SIG grant (#S10 OD026929). Computational analyses were performed using the San Diego Supercomputer Center’s Triton Shared Computing Cluster. This work was funded by the United States Government, Department of Energy, Advanced Research Projects Agency— Energy grant ARPA-E DE-FOA-0001858 to R.S.N. and L.M.L.L. with contracts to C.E.N., L.W.K., and E.E.A., National Science Foundation grants OCE-1837116 and EF-2025217 to E.E.A., and National Institutes of Health NIEHS grant R01-ES030316 to E.E.A.

