## Supplemental Figures S1, S2, S3 for "Enrichable consortia of microbial symbionts degrade macroalgal polysaccharides in *Kyphosus* fish"

**Supplementary Figure S1. Recovered MAGs from fish gut and enrichment metagenomes.** Enrichment sample names begin with the letter R, fish gut samples begin with the letter F. Fish gut samples were taken either from the midgut (GI) or hindgut (HG).

**Supplementary Figure S2. Complete gene tree with all binned GH86 CAZymes.** Phylogenetic tree with gene names colored by genome taxonomy. Cells mark the source of each CAZyme and whether SignalP predicts the presence of a signal peptide.

**Supplementary Figure S3. Motif logo of previously undescribed residue pattern for GH86-associated domain.** Logo created using WebLogo. Residue numbers are based on position in the pattern, rather than the parent protein. Few residues are conserved across all representatives.

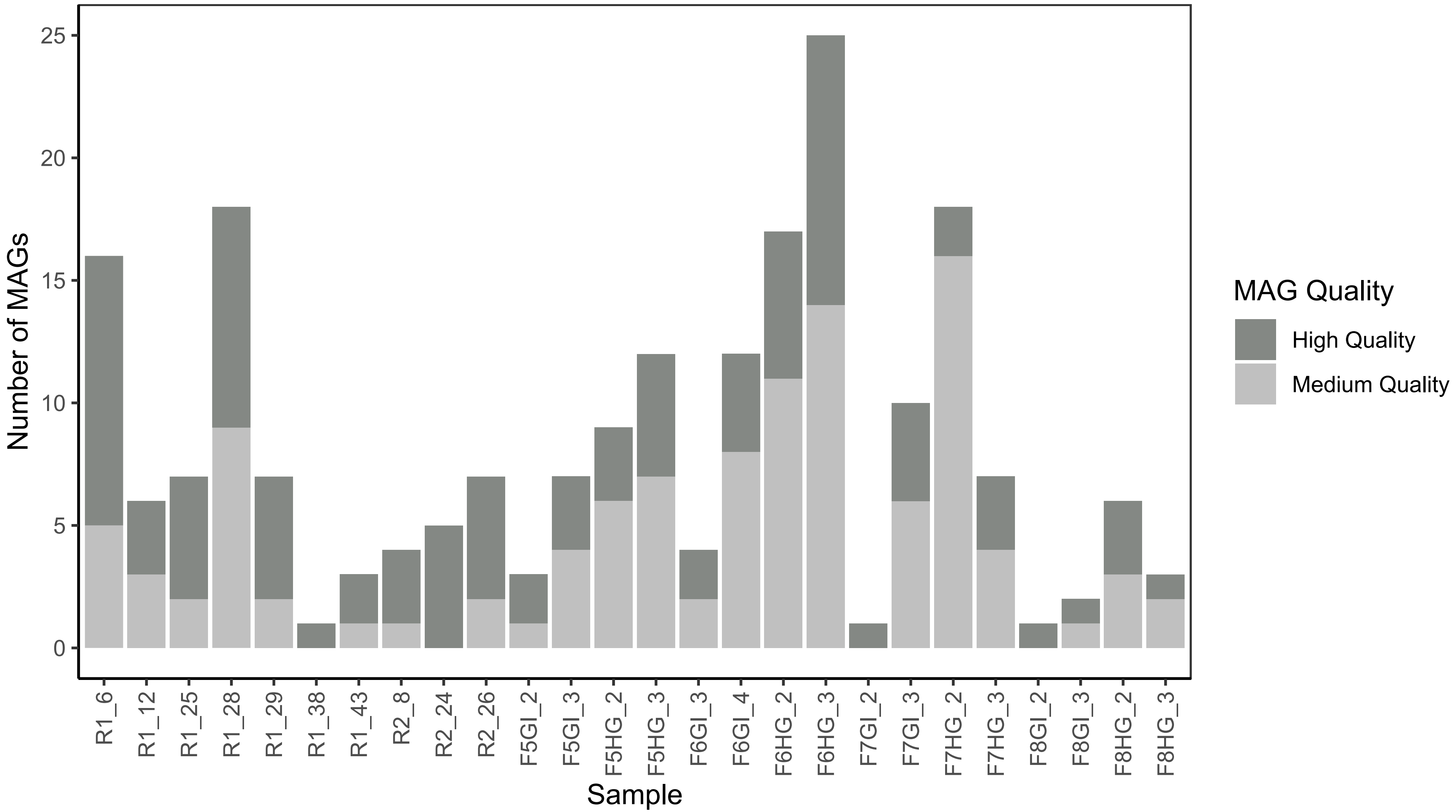


**Supplementary Figure S1. Recovered MAGs from fish gut and enrichment metagenomes.** Enrichment sample names begin with the letter R, fish gut samples begin with the letter F. Fish gut samples were taken either from the midgut (GI) or hindgut (HG).

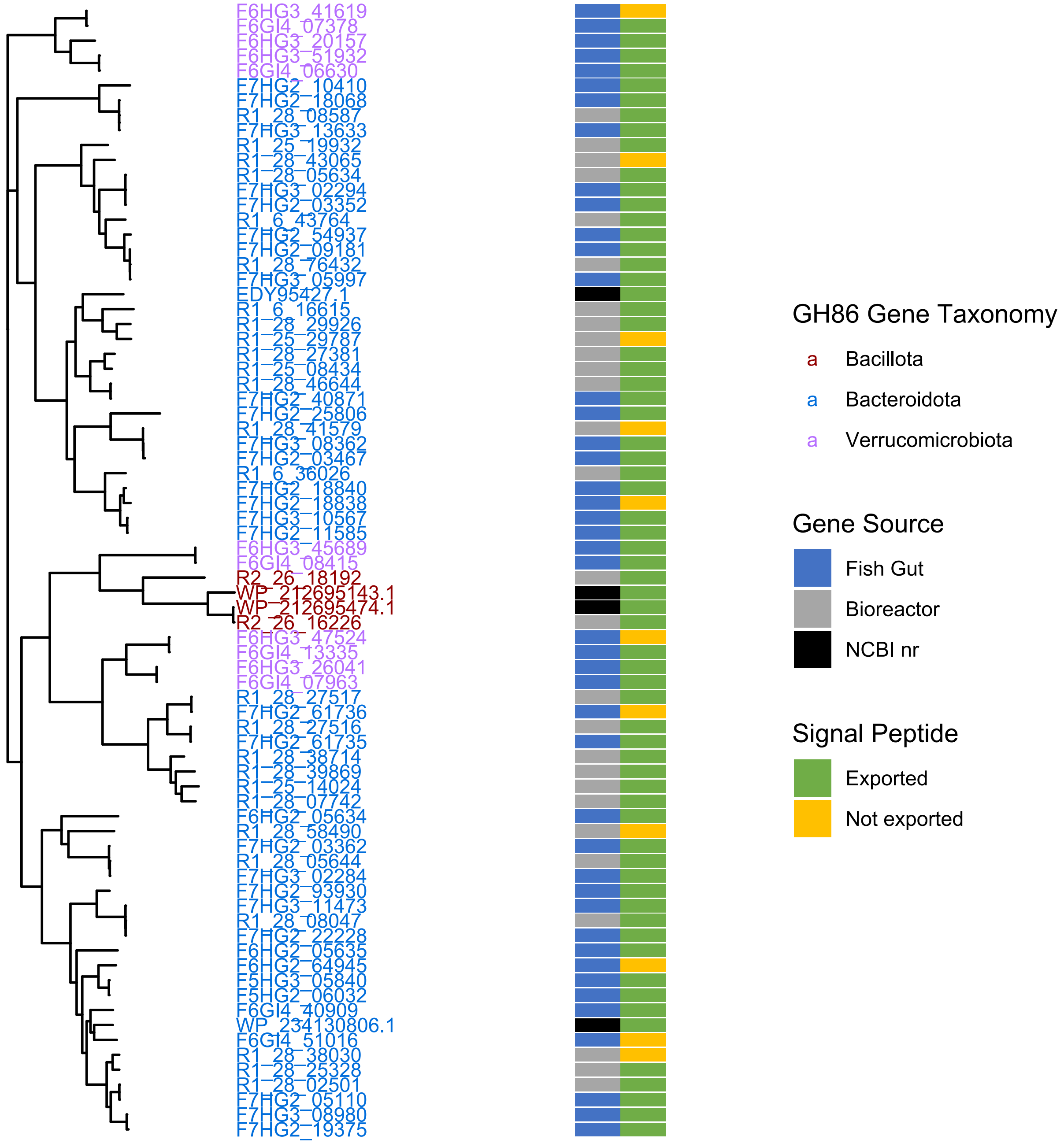


**Supplementary Figure S2. Complete gene tree with all binned GH86 CAZymes.** Phylogenetic tree with gene names colored by genome taxonomy. Cells mark the source of each CAZyme and whether SignalP predicts the presence of a signal peptide.

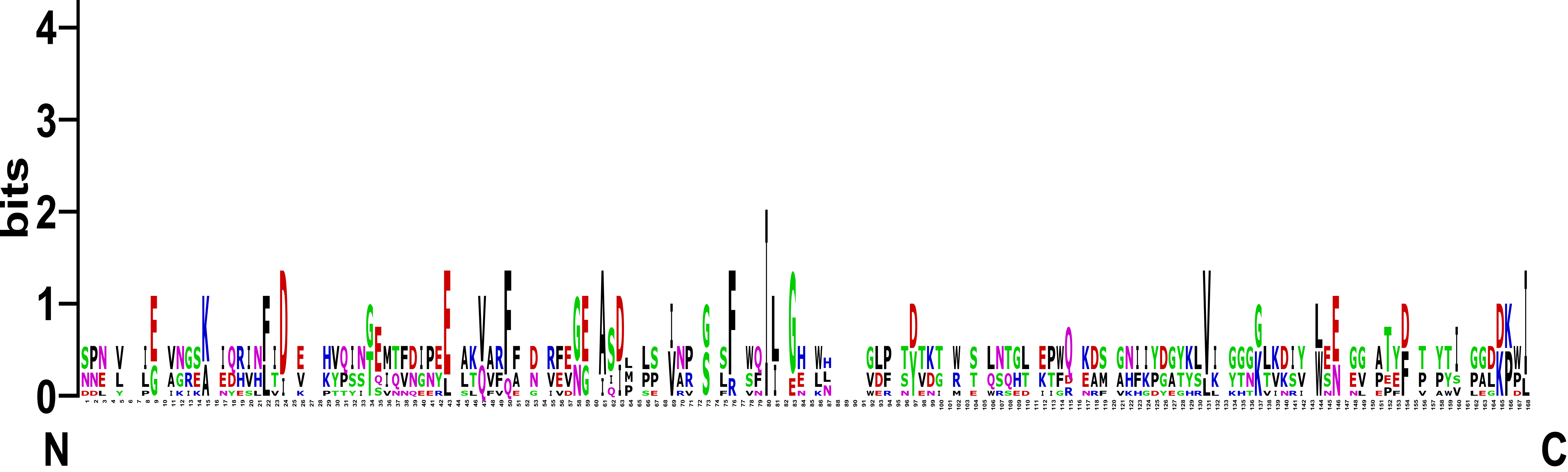


**Supplementary Figure S3. Motif logo of previously undescribed residue pattern for GH86-associated domain.** Logo created using WebLogo. Residue numbers are based on position in the pattern, rather than the parent protein. Few residues are conserved across all representatives.
